# Decorin enhances metabolic maturation by activating AMPK-PGC1A pathway in cardiac organoids

**DOI:** 10.1101/2024.06.20.599970

**Authors:** Myeong-Hwa Song, Seongmin Jun, Seung-Cheol Choi, Ji Eun Na, Im Joo Rhyu, Sun Wook Hwang, Minji Jeon, Do-Sun Lim

**Affiliations:** Department of Cardiology, Cardiovascular Center, College of Medicine, Korea University, 145 Anam-ro, Seongbuk-gu, Seoul, 02841, South Korea; R&D Center for Companion Diagnostic, SOL Bio Corporation, Suite 510, 27, Seongsui-ro7-gil, Seongdong-gu, Seoul 04780, Republic of Korea; Department of Anatomy College of Medicine, Korea University, 73, Goryeodae-ro, Seongbuk-gu, Seoul 02841, Republic of Korea; BK21 Graduate Program, Department of Biomedical Sciences, Korea University College of Medicine, Seoul, 02841, Republic of Korea; Department of Physiology, Korea University College of Medicine, Seoul, 02841, Republic of Korea; Department of Biomedical Informatics, Korea University College of Medicine, Seoul 02708, South Korea

## Abstract

**Rationale:** Cardiac organoids (COs) are advanced models for investigating heart development and disease, while require maturation to resemble the structural and functional characteristics of the human heart.

**Objective:** This study reveals the role of Decorin (DCN) contributes to the mature and vascularized COs and assesses the biological mechanism responsible for CO maturation.

**Methods and Results:** DCN-treated COs exhibit structural maturation involving aligned sarcomere, mitochondria, and t-tubule structures, and vessel formations, as well as functional maturation involving synchronized contraction-relaxation, Ca2+ transient, and increases ion channel expressions. DCN-treated COs also show metabolic maturation, including enhanced fatty acid oxidation and increased mitophagy. Transcriptional profiling results indicate that DCN-treated COs have increased levels of AMPK signaling and mitophagy. In DCN-treated COs, AMPK knockdown affects mitochondrial biogenesis, cardiac metabolism, ion channels, and mitophagy.

**Conclusions:** These findings indicate that DCN is crucial for development of mature, vascularized COs and that CO maturation is primarily regulated through AMPK signaling, which is triggered by DCN.

GRAPHIC ABSTRACT
A graphic abstract is available for this article.
DCN enhances metabolic maturation in COs by AMPK-triggered regulation of the glycolysis, fatty acid oxidation, and mitophagy.

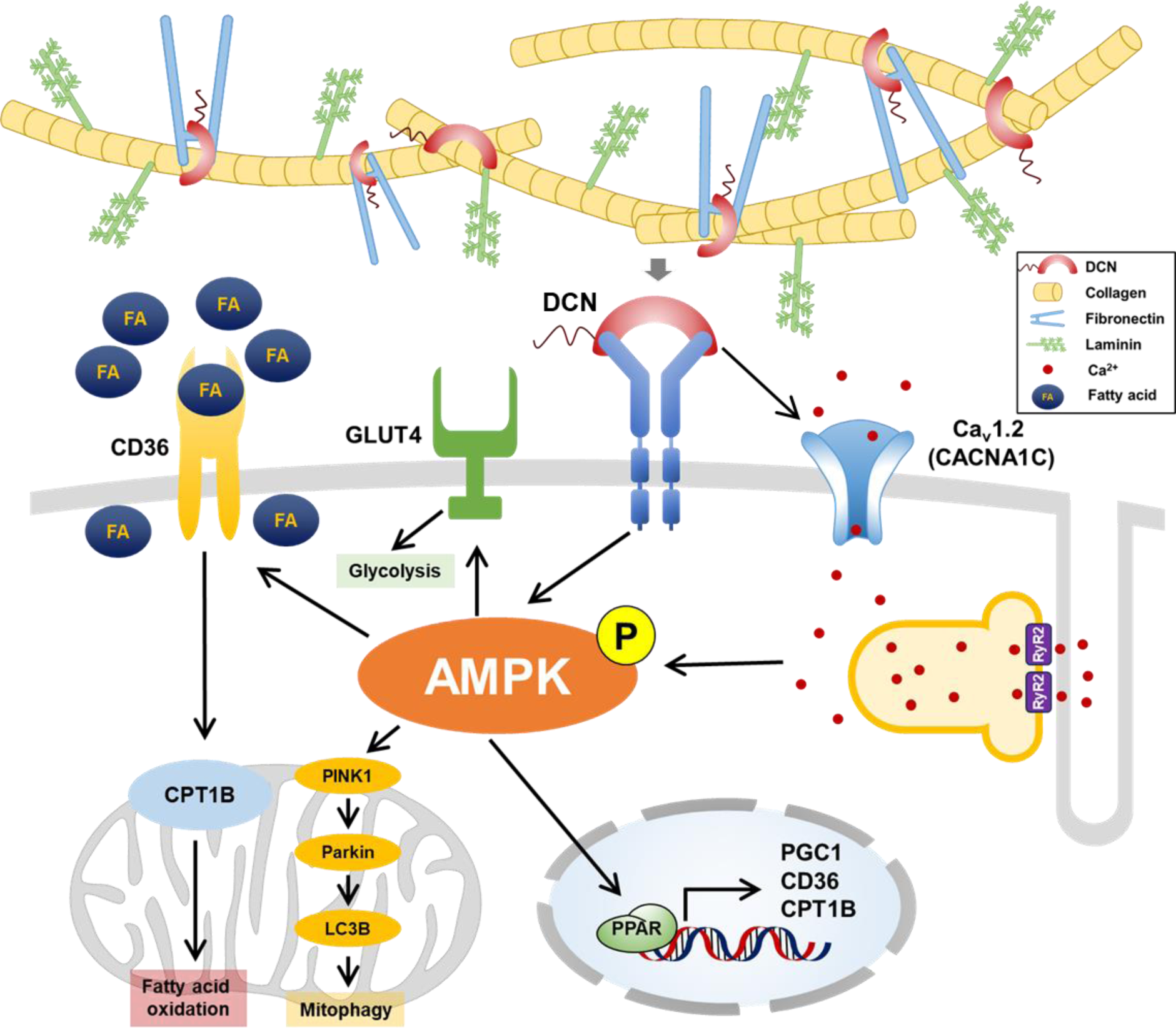

## INTRODUCTION

As cardiovascular disease (CVD) remains a top cause of mortality and morbidity worldwide, it is essential to pursue ongoing research into its mechanisms and develop effective treatment strategies. At heart of CVD lies dysfunction of cardiomyocytes (CMs), fundamental contractile cells responsible for maintaining proper cardiac function. Development and maturation of CMs are crucial processes that are essential to structural and functional integrity of heart. Thus, understanding mechanisms driving CM maturation is critical for advancing our knowledge of cardiac development and disease.

Differentiation of CMs using 3D culture techniques can result in mature CMs and imitate *in vivo* development process, encompassing aspects such as microenvironment, cell morphology, gene expression, and intracellular signal transduction^1–3^. Many studies using CMs differentiated human pluripotent stem cells (hPSCs) have leveraged these models for cardiac tissue simulation, disease modeling, and drug screening^4^. However, achieving differentiation into fully mature CMs remains a pivotal challenge; thus, developing technologies that can efficiently induce such maturation is of paramount importance.

CM maturation involves progression from an immature, feta-like state to a more mature, adult-like state, encompassing structural, functional, and metabolic enhancements^5,6^. First, Structural maturation of CMs is involved in organized sarcomeres and mitochondria and longer sarcomere length and Z-line width^7,8^. Structurally mature CMs also exhibited mature transverse-tubules (t-tubules)^9,10^. Second, functionally mature CMs show prolonged action potential duration, contraction-relaxation duration, peak-to-peak duration, and increased Ca^2+^ handling by RyR-mediated Ca^2+^ release^3,11–13^. Lastly, metabolically matured CMs showed increased cardiac metabolism and ATP production for contraction and relaxation, and increased lipid substrates for promoting metabolism by fatty acid oxidation (FAO)^10,14–16^.

Cardiac organoids (COs), developed in 3D culture systems, include cardiac structure and function, and comprise various cardiac component cells such as CM, endothelial cells (ECs), and fibroblasts (FB)^17^. Therefore, COs replicate stiffness, tissue density, and reactivity of cardiac tissue. Growth factors and mechanically generated stresses significantly influence heart structure development, with a crucial role played by concentration gradient of these factors^18^. Additionally, vascular network formation within organoids is vital for efficient exchange of nutrients and waste and for establishing self-organizing features such as angiogenesis and permeability^19,20^. In this study, cardiac component cells (CMs, ECs, and FBs) were analyzed to investigate COs similar to heart, consisting of various cells, through overexpression of target gene on CO maturation.

Decorin (DCN) is an extracellular matrix (ECM) protein and proteoglycan that plays important roles in cellular processes by interacting with collagen and fibronectin to enhance tissue integrity^21^. DCN promotes PINK1/Parkin-mediated mitophagy pathway through PGC1A and Mitostatin activation and supports autophagy via AMPK phosphorylation along with Beclin1 and LC3 activation^22^. In injured hearts, DCN can inhibit excessive mitophagy by modulating PGC1A-Mitostatin pathway^23,24^. Furthermore, DCN regulates various receptors including IGF1R, EGFR, and TGFβ, which are instrumental in inducing CM maturation^25–27^. By binding to TGFβ, DCN triggers signaling pathways that promote transcription and inhibits TGFβ-SMAD2/3 pathway to mitigate inflammation and fibrosis, thereby reducing cardiac remodeling and dysfunction in CVD^23^. Additionally, DCN is interaction with VEGFR2 facilitates angiogenesis and mitophagy.

This study demonstrates the effects of DCN overexpression in COs to analyze maturation processes. We have established that DCN significantly enhances maturation of both general and ventricular CMs. Moreover, using transcriptome analysis and gene knockdown approaches, we identified specific genes and pathways that facilitate development of mature and structurally integrated ventricular COs.

## METHODS

For detailed methodological information, please refer to the Data Supplement.

## RESULTS

Through RNA-seq analysis of matured COs, DCN was identified as a factor that can induce maturation of COs. To evaluate its effect on maturation, DCN was administered from differentiation day 5 to day 30 (Figure 1A). Phase-contrast images showed no significant differences in morphology (Figure 1B). The CM subtypes in both control and DCN-treated COs was assessed by qRT-PCR for markers of CM subtype on day 30 (Figure 1C). mRNA level of *MLC2v* was notably higher in DCN-treated COs. However, mRNA levels of *MLC2a* and *TBX18* were not significantly differed between control and DCN-treated COs. Flow cytometry analysis further investigated proportions of CM subtypes (Figure 1D), revealing a higher percentage of MLC2v (38.1% vs 52.5%) was higher in DCN-treated COs compared with control, while the percentages of MLC2a (7.31% vs 9.00%) and TBX18 (1.20% vs 0.88%) were higher in control compared with DCN-treated COs. Furthermore, protein level of MLC2v was increased in DCN-treated COs relative to control, while protein levels of MLC2a and TBX18 showed not significantly differences between the groups (Figure 1E). Subsequently, immunofluorescence staining was used to analyze the proportion of cTnT+ and MLC2v+ CMs, as well as MLC2a+ and MLC2v+ CMs, in both the control and DCN-treated COs (Figure 1F and 1G). The results showed a higher proportion of MLC2v+ CMs in the DCN-treated COs compared with control. These findings revealed that DCN-treated COs predominantly consist of ventricular-like CMs compared to control.

**Figure 1.**
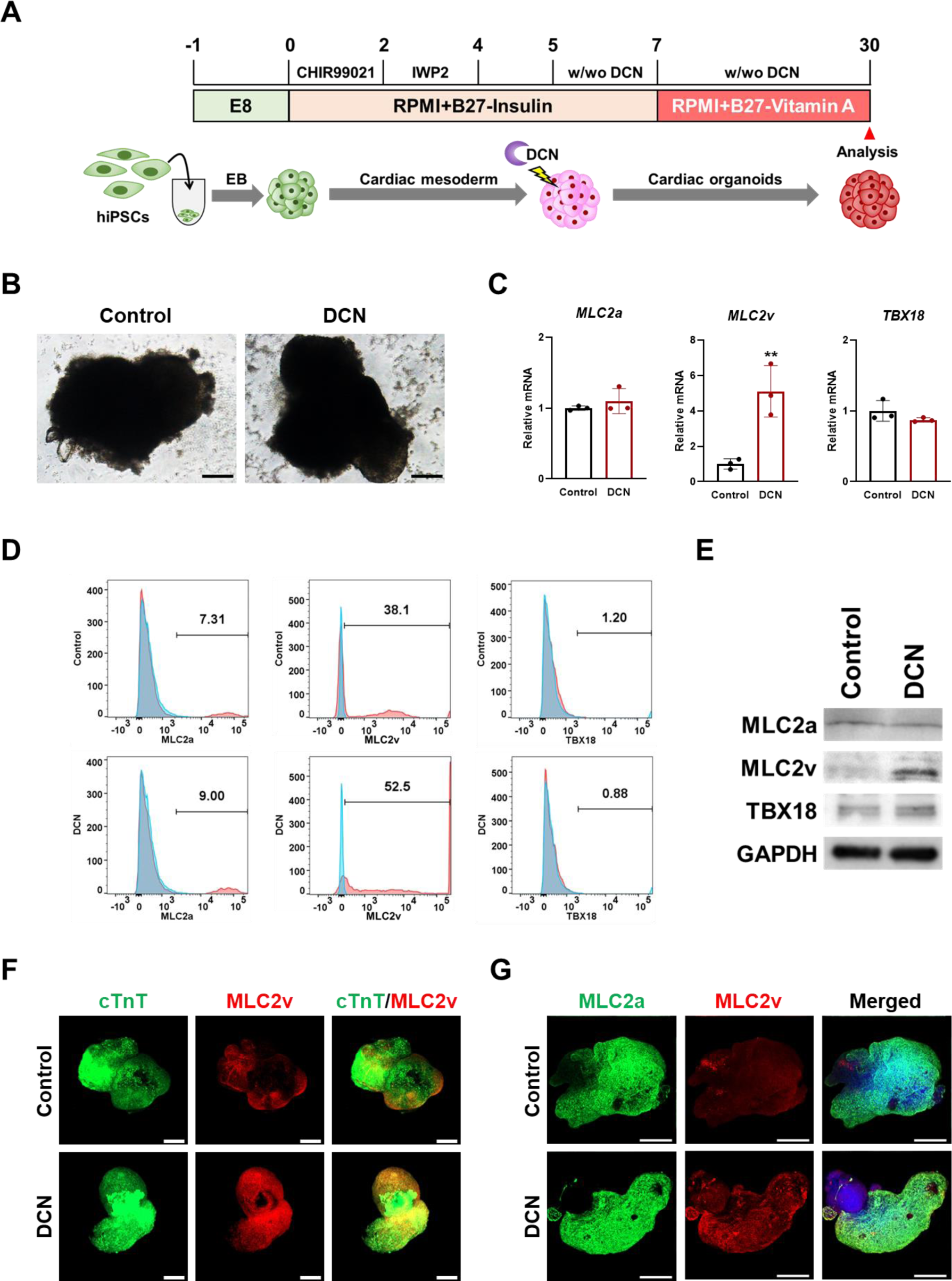
DCN-treated COs were predominantly composed of ventricular CMs. **A,** Schematic diagram of CO formation with/without DCN. **B,** Phase-contrast images of control and DCN-treated COs. Scale bars = 200 μm. **C,** qRT-PCR of atrial (*MLC2a*), ventricular (*MLC2v*), and nodal (*TBX18*) CM markers in control and DCN-treated COs. n=3 for each group. Values represent means ± SDs. **p<0.01. **D,** Flow cytometry analysis of MLC2a, MLC2v, and TBX18 in control and DCN-treated COs. **E,** Western blot analysis of MLC2a, MLC2v, and TBX18 in control and DCN-treated COs, with GAPDH as a loading control. **F,** Immunofluorescence analysis of cTnT (green) and MLC2v (red) in control and DCN-treated COs. Nuclei stained with DAPI (blue). Scale bars = 200 μm.

### CD31+ ECs were found in abundance in DCN-treated COs

Next, mRNA expression levels of markers for cardiac component cells, such as CMs, ECs, and FBs, were analyzed using qRT-PCR (Figure 2A). Expression levels of *cTnT* and *CD31* (EC marker) were significantly elevated in DCN-treated COs compared with control, while expression of *FSP1* (FB marker) was no significant difference in control and DCN-treated COs. Consequently, flow cytometry analysis was conducted to identify the proportion of cardiac component cells in COs (Figure 2B). The percentages of cTnT (40.7% vs 53.8%) and CD31 (14.4% vs 27.4%) were notably higher in DCN-treated COs compared with control, whereas the percentage of FSP1 (20.1% vs 12.1%) was similar to control and DCN-treated COs. Protein expressions of cTnT and CD31 were elevated in DCN-treated COs compared with control, while protein level of FSP1 was not significantly different between control and DCN-treated COs.

**Figure 2.**
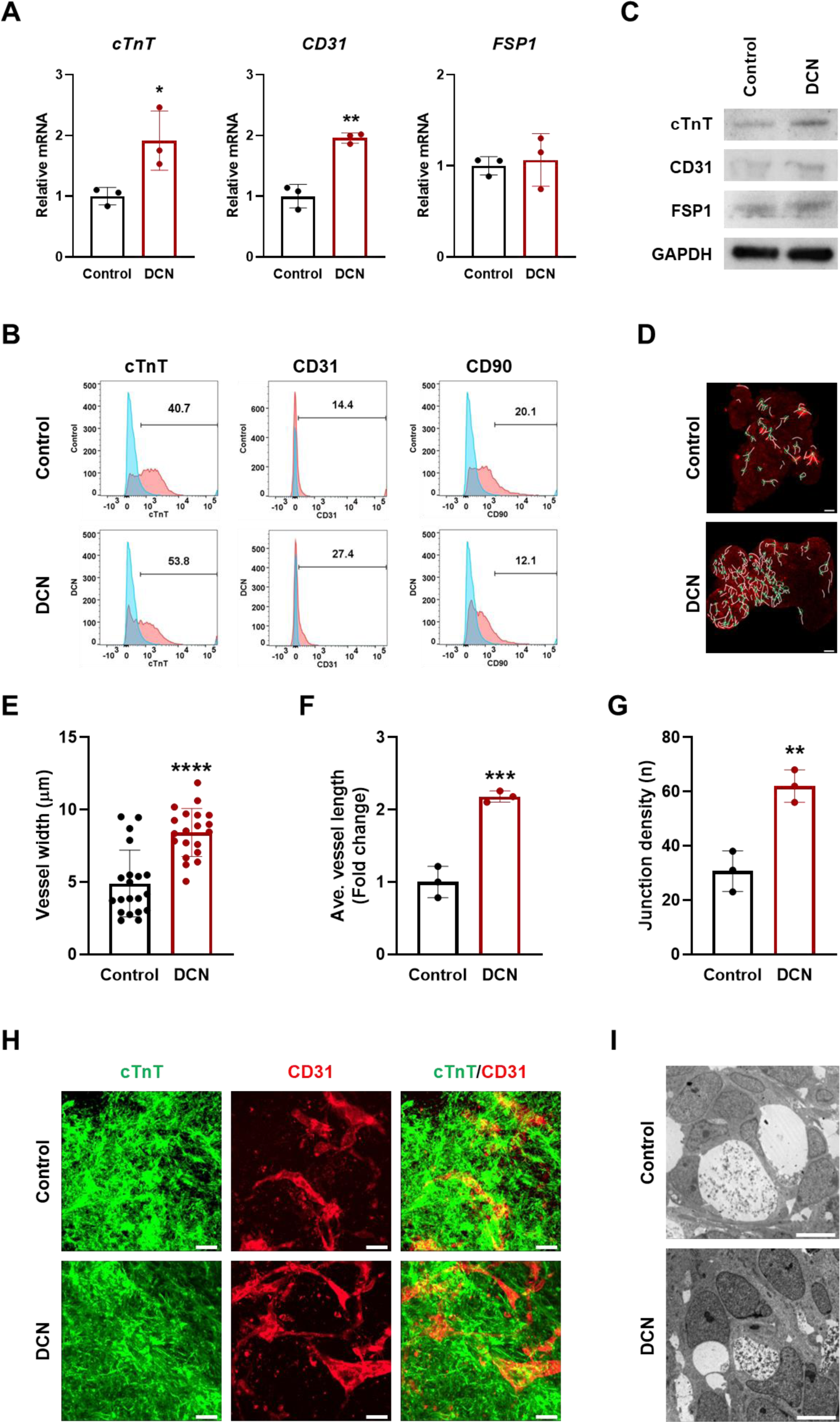
DCN improved vessel width, length, and large vessels in COs. **A,** qRT-PCR of genes encoding CM (*cTnT*), EC (*CD31*), and FB (*FSP1*) markers in control and DCN-treated COs. Values represent means ± SDs. n = 3 for each group. *p < 0.05, **p < 0.01. **B,** Western blot analysis of cTnT, CD31, and FSP1 in control and DCN-treated COs, with GAPDH as a loading control. **C,** Flow cytometry analysis of cTnT, CD31, and FSP1 in control and DCN-treated COs. **D,** Immunofluorescence images showing CD31+ EC (red). White lines: vessels, green circles: junctions. Scale bars = 50 μm. Graphical representation of **E,** vessel width, **F,** average vessel length, and **G,** junction density. Values represent means ± SDs. n=20 for vessel width and n=3 for average vessel length and junction density. *p < 0.05, **p < 0.01, ***p < 0.001. **H,** Immunofluorescence images showing cTnT+ CMs (green) and CD31+ ECs (red). Nuclei were stained with DAPI (blue). Scale bars = 20 μm. **I,** TEM images showing the ECs and vascular lumens in control and DCN-treated COs. Scale bars = 2 μm. vascular lumens (L). The dotted boxes indicated vessels.

To examine vascular structure, the expression of CD31 was assessed using immunofluorescence staining (Figure 2D). The vessel width, average vessel length, and junction density were analyzed based on CD31-stained images (Figure 2E-G). Immunofluorescent staining was used to visualize both CMs and vascular structures in COs, revealing the presence of cTnT and CD31 in both control and DCN-treated COs. However, CD31+ vessels were more prevalent in DCN-treated COs (Figure 2H). Therefore, TEM was employed to examine vessel structure, showing a greater number of lumens composed of two or more ECs in DCN-treated COs compared with control (Figure 2I). These results indicate that larger vessels were present in DCN-treated COs.

### DCN-treated COs are structurally matured compared with control

Adult CMs are characterized by aligned sarcomeres and mitochondria, the presence of t-tubule structures, larger sizes, and elongated mitochondria^28,29^. Furthermore, adult CMs possess longer sarcomere length and greater number of mitochondrial compared to immature CMs. Structural maturity COs was assessed by examining sarcomeres through immunofluorescence staining and TEM (Figure 3A and 3B). Organized sarcomeres, indicative of maturation, were more prevalent in DCN-treated COs than control, with sarcomere lengths averaging 2.1 μm in DCN-treated COs and 1.8 μm in control (Figure 3C). Moreover, Z-line width was longer in DCN-treated COs compared with control (Figure 3D). Cardiac t-tubules, invaginations of CM sarcolemma, are crucial for excitation-contraction coupling and action potential regulation due to association with numerous ion channels^30^. T-tubule formation was investigated through qRT-PCR analysis of t-tubule markers (Figure 3E), revealing significantly higher expression levels of *CAV3*, *JPH2*, and *BIN1* in DCN-treated COs. Protein levels of t-tubule markers were further confirmed by western blot analysis, with CAV3 and JPH2 showing pronounced expression in DCN-treated COs (Figure 3F). Immunofluorescence staining was utilized to visualize t-tubule formation and co-localization for t-tubule in CMs (Figure 3G), demonstrating an abundance of JPH2+ CMs and a higher expression of co-localized cTnT+ and JPH2+ CMs in DCN-treated COs compared with control.

**Figure 3.**
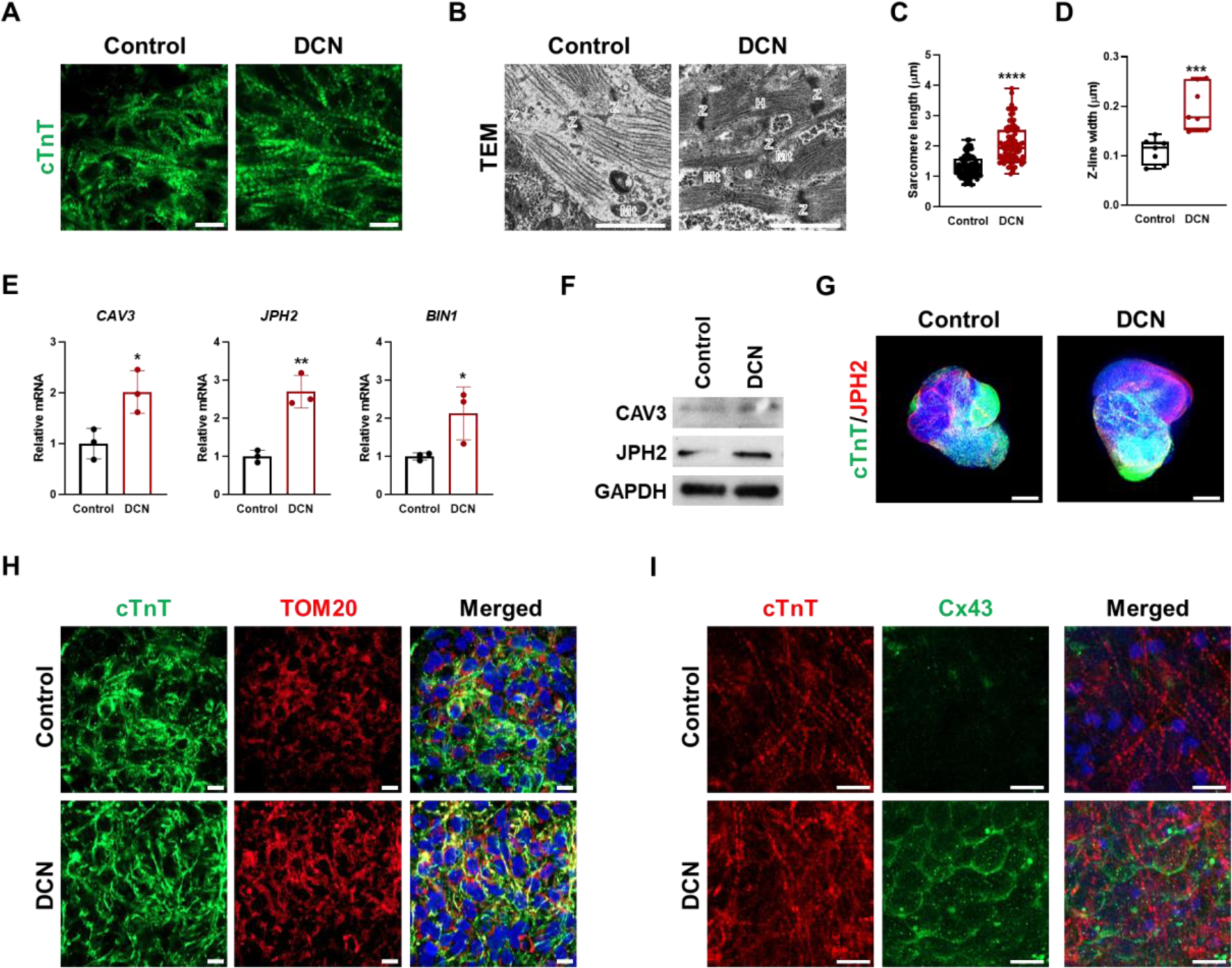
DCN-treated COs exhibit structural maturation. **A,** Fluorescence analysis of cTnT (green) in control and DCN-treated COs. Scale bars = 10 μm. **B,** TEM images showing the sarcomere in control and DCN-treated COs. Scale bars = 1 μm. Quantification of **C,** sarcomere length and **D,** Z-line width in control and DCN-treated COs. Values represent means ± SDs. n = 12 for each group. *p < 0.05. **E,** qRT-PCR of t-tubule markers (*CAV3*, *JPH2*, and *BIN1*) in control and DCN-treated COs. Values represent means ± SDs. n = 3 for each group. *p < 0.05, **p < 0.01. **F,** Western blot analysis of CAV3 and JPH2 in control and DCN-treated COs. GAPDH served as a loading control. **G,** Immunofluorescence analysis of cTnT (green) and JPH2 (red) in control and DCN-treated COs. Nuclei were stained with DAPI (blue). Scale bars = 200 μm. Immunofluorescence analysis of **H,** cTnT (green) and a mitochondria marker, TOM20 (red), and **I,** cTnT (green) and gap junction marker, Cx43 (red), in control and DCN-treated COs. Nuclei were stained with DAPI (blue). Scale bars = 10 μm.

Organized sarcomeres and mitochondrial alignment were observed in mature CMs^8,31^. Therefore, Sarcomere and mitochondrial alignment were investigated using immunofluorescence staining (Figure 3H). Co-localized areas of cTnT and a mitochondrial marker, TOM20, showed higher expression in DCN-treated COs compared with control. Connexin 43 (Cx43), a predominant gap junction protein in the ventricular myocardium, plays a crucial role in cardiac excitation and contraction^32^. Immunofluorescence staining revealed that in DCN-treated COs, Cx43 was uniformly expressed at cell junctions, whereas in the control group, Cx43 displayed a partial distribution (Figure 3I).These results revealed that DCN treatment promotes structural maturation in COs.

### The beating property of DCN-treated COs represent mature CMs

To analyze functional maturation in COs, beating properties of both control and DCN-treated COs were examined using movies captured by a microscope (Figure 4A and Movie 1 and 2). Analysis of the difference between contraction and relaxation phases revealed that DCN-treated COs had a greater contraction-relaxation interval compared to the control, indicating stronger beating activity (Figure 4A). The frequency of beats per minute (BPM) was significantly lower in DCN-treated COs compared with control, whereas both peak-to-peak duration and contraction-relaxation duration were increased (Figure 4B-D). In adult CMs, Ca^2+^-handling proteins are critical for excitation-contraction coupling, and levels of L-type Ca^2+^ channels are higher in ventricular CMs than in immature CMs^33,34^. Given that calcium transients are essential for excitation-contraction coupling in CMs, concentration of Ca^2+^ was measured using specific channel markers in both control and DCN-treated COs (Figure 4E). mRNA expression levels of Ca^2+^ channel markers (*RyR2* and *CACNA1C*) were significantly increased in DCN-treated COs. Furthermore, Fluo-4 AM fluorescence intensity, which is synchronized with contraction-relaxation cycles, was stronger in DCN-treated COs compared with control organoids (Figure 4F and I).

**Figure 4.**
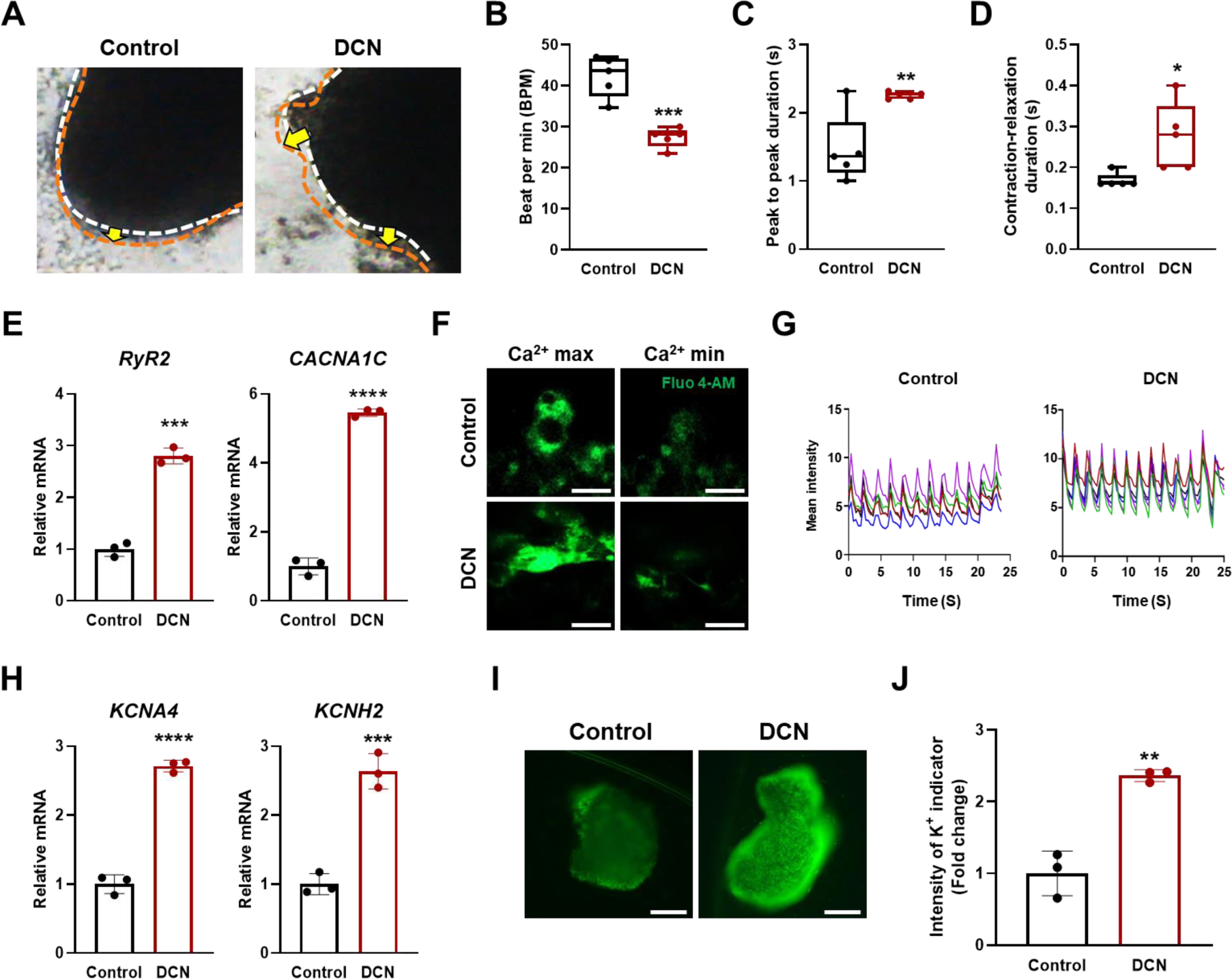
DCN induces functional maturation involved in spontaneous contractility and ion channel activity. **A,** Images contraction (White)-relaxation (Orange) in COs. Quantification of **B,** BPM, **C,** peak-to-peak duration, **D,** contraction-relaxation duration in control and DCN-treated COs. Values are means ± SDs. n=5 for each group. **p < 0.01, ***p < 0.0001. **E,** qRT-PCR of calcium channel markers (*RyR2* and *CACNA1C*) in control and DCN-treated COs. Values are means ± SDs. n=3 for each group. ***p < 0.001, ****p < 0.001. **F,** Fluorescence images showing maximum (Ca2+ max) and minimum (Ca2+ min) intensity of Fluo-4AM+ cells (green). **G,** Graph showing intensity of Fluo-4 AM in control and DCN-treated COs. **H,** qRT-PCR of potassium channel markers (*KCNA4*, *KCNH2*, and *KCNJ2*) in control and DCN-treated COs. Values are means ± SDs. n=3 for each group. ***p < 0.0001. **I,** Immunofluorescence images showing COs stained with potassium indicator.

The action potential of CMs is influenced by various ions and ion channels, with ion channel-related genes being upregulated in sarcolemma of adult CMs^35^. Therefore, the expression of genes encoding potassium channel markers (*KCNA4*, *KCNH2*, and *KCNJ2*) were investigated using qRT-PCR (Figure 4H). mRNA levels of *KCNA4*, *KCNH2*, and *KCNJ2* were higher in DCN-treated COs than in control. Potassium channel function was further assessed using specific indicators, showing an increase in potassium indicators in DCN-treated COs (Figure 4I and 4J). These finding indicate that DCN treatment promotes functional mutation of COs.

### DCN induces metabolic maturation in COs

To analyze metabolic maturation in COs, mitochondrial maturity was initially assessed through TEM images (Figure 5A). DCN-treated COs showed complex and elongated mitochondria, in contrast to small and round mitochondria observed in control. Both mitochondrial number and cristae densities were significantly higher in DCN-treated COs compared with control (Figure 5B and C). To quantify mitochondrial content in control and DCN-treated COs, pRT-PCR was performed using mitochondrial DNA (mtDNA) and nDNA. Ratio of mtDNA to nDNA was notably higher in DCN-treated COs compared with control, suggesting an increased mitochondrial content in DCN-treated COs.

**Figure 5.**
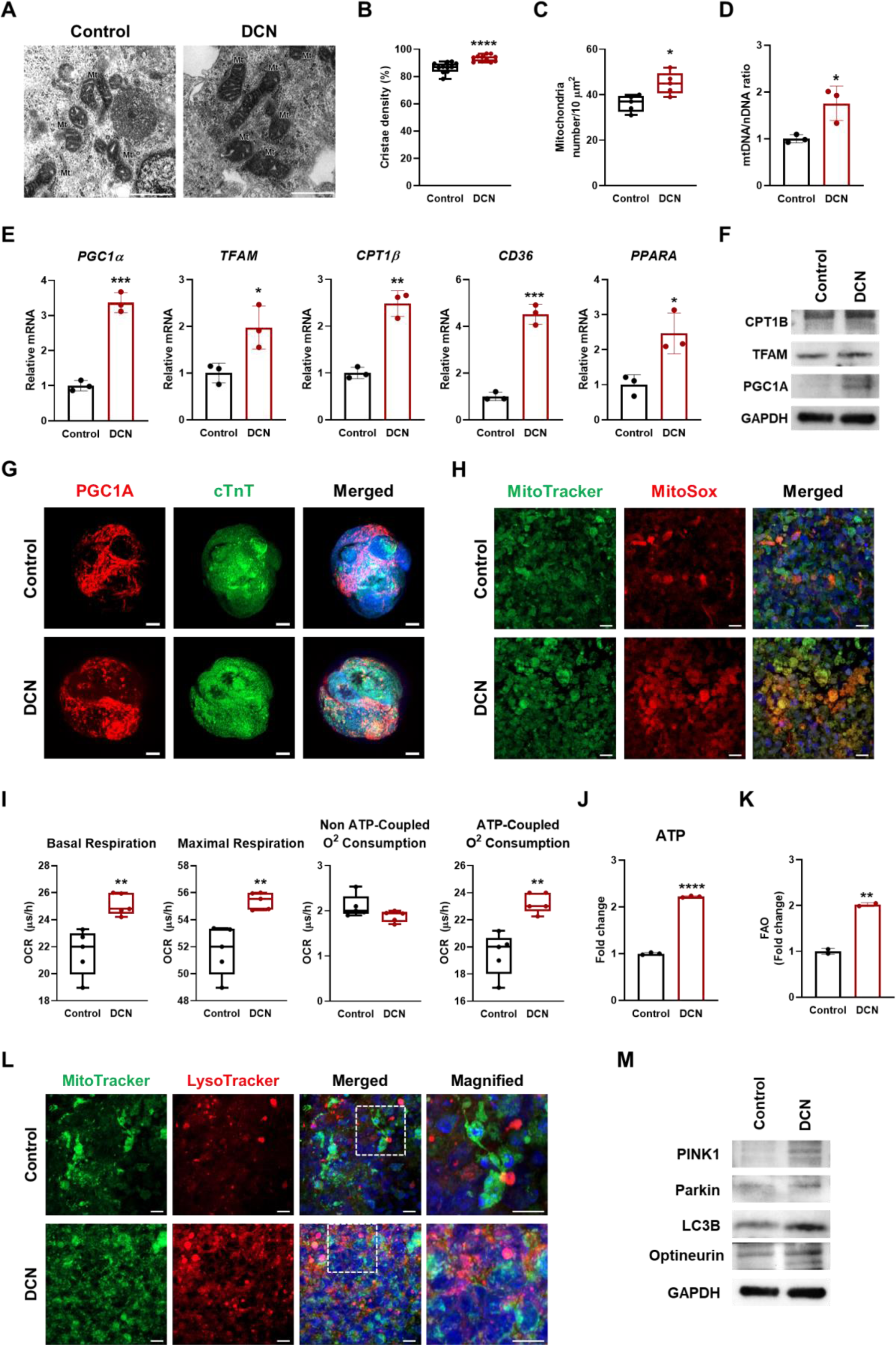
DCN enhances mitochondrial maturation and cardiac metabolism in COs. **A,** TEM images showing the mitochondria in control and DCN-treated COs. Scale bars = 1 μm. Quantification of **B,** mitochondrial numbers and **C,** cristae densities in control and DCN-treated COs. Values represent means ± SDs. n = 5 for B and n = 11 for C. *p < 0.05, ****p < 0.0001. qRT-PCR analysis showing **D,** ratio of mtDNA/nDNA and **E,** expression of cardiac metabolic markers (*PGC1*α, *TFAM*, *CPT1*β, *CD36*, and *PPARA*) in control and DCN-treated COs. Values represent means ± SDs. n = 3 for each group. *p < 0.05, **p < 0.01, ***p < 0.0001. **F,** Western blot analysis showing CPT1B, TFAM, and PGC1A in control and DCN-treated COs with GAPDH as a loading control. Immunofluorescence images showing **G,** cTnT (green), PGC1A (red), **H,** MitoTracker (green), and MitoSox (red) in control and DCN-treated COs. Nuclei were stained with DAPI (blue). Scale bars = 10 μm. OCR analysis showing **I,** basal respiration, **J,** Maximal respiration, **K,** non-ATP-coupled O2 consumption rate, and **L,** ATP-coupled O2 consumption rate in control and DCN-treated COs. Values represent means ± SDs. n = 5 for each group. **p < 0.01. **M,** ATP assay showing ATP contents in control and DCN-treated COs. Values represent means ± SDs. n = 3 for each group. ****p < 0.0001.

Genes related to mitochondrial biogenesis (*PGC1A* and *TFAM*) and FAO (*CPT1B*, *CD36*, and *PPARA*) were quantified by qRT-PCR (Figure 5E). DCN-treated COs exhibited significantly higher mRNA levels of *PGC1A*, *TFAM*, *CPT1B*, *CD36*, and *PPARA* compared with control. Similarly, protein levels of PGC1A, TFAM, and CPT1B were increased in DCN-treated COs relative to control (Figure 5F). Immunofluorescence staining revealed that the expression of mitochondrial biogenesis marker (PGC1A) was substantially higher in DCN-treated COs compared with control.

Given that reactive oxygen species (ROS) can activate cardiac differentiation and maturation through various extracellular signals, mitochondrial ROS was analyzed using MitoTracker, a total live mitochondria indicator, and MitoSox, which indicates ROS produced by mitochondria^36,37^. Co-localization of MitoTracker and MitoSOX was greater in DCN-treated COs than in control (Figure 5H), suggesting that DCN promotes mitochondrial maturation.

Mitochondrial function and respiratory reserve capacity were investigated using OCR analysis. Cardiac metabolism was analyzed for metabolic maturation by measuring OCRs (Figure 5I). Basal respiration, maximal respiration, and ATP-coupled O2 consumption rates were higher in DCN-treated COs than in control, while non-ATP-coupled O2 consumption rate showed no significant difference between control and DCN-treated COs. ATP content in COs was quantified using an ATP assay (Figure 5J), revealing that ATP content was predominantly higher in DCN-treated COs compared to control. Since mature CMs obtain most energy through FAO^38^, an investigation into FAO revealed a significant increase in FAO in DCN-treated COs (Figure 5K). These results demonstrate that DCN treatment leads to metabolic maturation in COs.

### Mitophagy was increased in DCN-treated COs compared with control

In CMs, mitochondrial maturation is induced by various factors, including metabolic signals. These metabolic signals play a crucial role in development and maturation of CMs^39^. Mitophagy, in particular, regulates mitochondrial biogenesis, maturation, metabolism, and differentiation in CM^40^. Activity of mitophagy in COs was investigated. Initially, to assess mitophagy in COs, co-localization of MitoTracker and LysoTracker was examined (Figure 5L). Areas showing co-localization of MitoTracker and LysoTracker were significantly more pronounced in DCN-treated COs compared with control.

Dysfunctional and damaged mitochondria are recycled through PINK1-Parkin-mediated mitophagy, which is crucial for maintaining mitochondrial quality and function^41^. Western blot analysis was conducted to examine mitophagy signaling pathway (Figure 5M). Protein expression levels of PINK1, Parkin, LC3B, and optineurin was found to be upregulated in DCN-treated COs compared with control. These findings suggest that DCN enhances mitochondrial functions, such as mitophagy, contributing to metabolic maturation of COs.

### Transcriptome analysis indicated an upregulation of cardiac-related genes in DCN-treated COs compared with control

To identify genes and mechanisms responsible for morphological and physiological maturation, transcriptome analysis was performed on both control and DCN-treated COs at day 30. Heatmap analysis indicated an upregulation of cardiac-specific genes in both control and DCN-treated COs, while other organ-specific genes were also noted (Figure 6A). Furthermore, heart-specific genes were found to be more highly expressed in DCN-treated COs compared with control. A scatter plot was constructed to highlight DEGs, particularly those related to heart, showing that these genes were expressed more highly in DCN-treated COs than in control(Figure 6B). A scatter plots further underscored genes that were upregulated in control versus DCN-treated COs, with upregulated genes predominantly related to cardiac function. A TRRUST Transcription Factors graph observed the heart development, cardiac metabolism, and endothelial cell related transcription factors upregulated in DCN-treated COs compared with control (Figure 6C).

**Figure 6.**
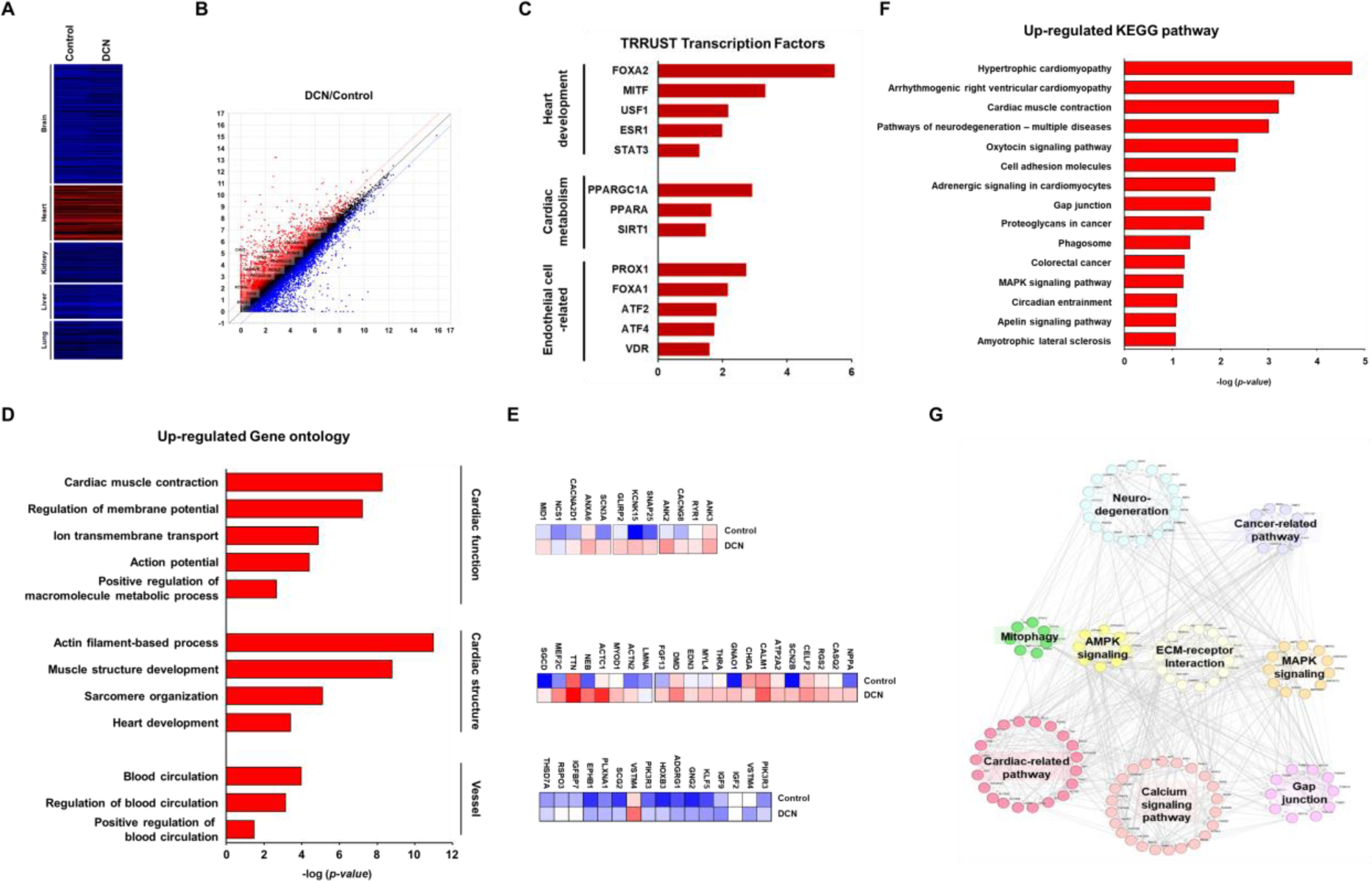
Transcriptome analysis indicates that genes involved in cardiac maturation and vessel formation are upregulated in DCN-treated COs compared with control. **A,** Heatmap of organ-specific genes obtained from RNA-seq. **B,** Scatter plot showing transcript expression in control and DCN-treated COs. **C,** TRRUST transcription factors for DEGs. **D,** GO analysis of upregulated genes in DCN-treated COs compared with control. **E,** Heatmap of cardiac function (top), cardiac structure (middle), and vessel (bottom) in control and DCN-treated COs. **F,** Pathway analysis of the major 15 enriched KEGG pathways in DCN-treated COs compared with control. **G,** Network analysis of genes highlighted in F according to translated protein-protein interactions using STRING database.

To evaluate biological implications of these gene expression changes, DEGs in DCN-treated COs compared with control were categorized using GO analysis (Figure 6D). The analysis revealed that the most significantly upregulated DEGs in DCN-treated COs were associated with cardiac function, cardiac structure, and angiogenesis. The most significant DEGs are presented in heatmaps in figure 6E.

### RNA-seq analysis shows that DCN-treated COs exhibit cardiac maturation via ECM-receptor, MAPK, and AMPK signaling pathway

To determine signaling pathways associated with DEGs in control and DCN-treated COs, transcriptomes were examined using KEGG pathway databases (Figure 6F). The top 15 KEGG pathways associated with DEGs in DCN-treated COs compared to control include those related to cardiomyopathies, cardiac muscle contraction, neurodegenerative diseases, cell adhesion, adrenergic signaling, gap junctions, cancer, phagosome, MAPK signaling, circadian entrainment, and apelin signaling. These comparative results from RNA-seq analysis of control and DCN-treated COs indicate that DCN treatment leads to structural, functional, and metabolic maturation, as well as vessel formation in COs (Figure 6G).

Potential mechanisms identified through transcriptional profiling, including ECM-receptor interaction, MAPK signaling pathway, and AMPK signaling, were validated using qRT-PCR and western blot analysis for maturation of DCN-treated COs. The ECM in heart is vital for supporting fundamental cellular processes during both organ development and homeostasis. Transmembrane heterodimeric receptors known as integrins are involved in key cellular activities, including cell-ECM adhesion, organization of cellular structure, and converting mechanical signals from ECM into biochemical signals in CMs.

mRNA expression levels of collagen (*COL1A1*, *COL3A1*, and *COL4A1*), fibronectin (*FN1*), and laminin (*LAMA2*) markers were significantly higher in DCN-treated COs compared with control (Figure 7A-C). Similarly, protein expressions of COL1A1, FN1, and Laminin were elevated in DCN-treated COs compared with control (Figure 7D). To investigate ECM downstream signaling, the levels of membrane protein complexes (SGCA, SGCB, SGCD, SGCG, DES, DMD) and Integrins were examined (Figure 7E-H). The mRNA levels of SGCA, SGCB, SGCD, SGCG, DES, DMD, and ITGA7 were elevated in DCN-treated COs. The protein levels of ITGA5, ITGB3, and ITGB4 were also increased in DCN-treated COs. In addition, levels of focal adhesion markers that are downstream signals of integrins were examined (Figure 7H). Expression levels of p-FAK, ROCK1, LIMK1, and p-cofilin were upregulated in DCN-treated COs compared with control (Figure 7I). These results suggest that DCN activates ECM–integrin–focal adhesion in COs.

**Figure 7.**
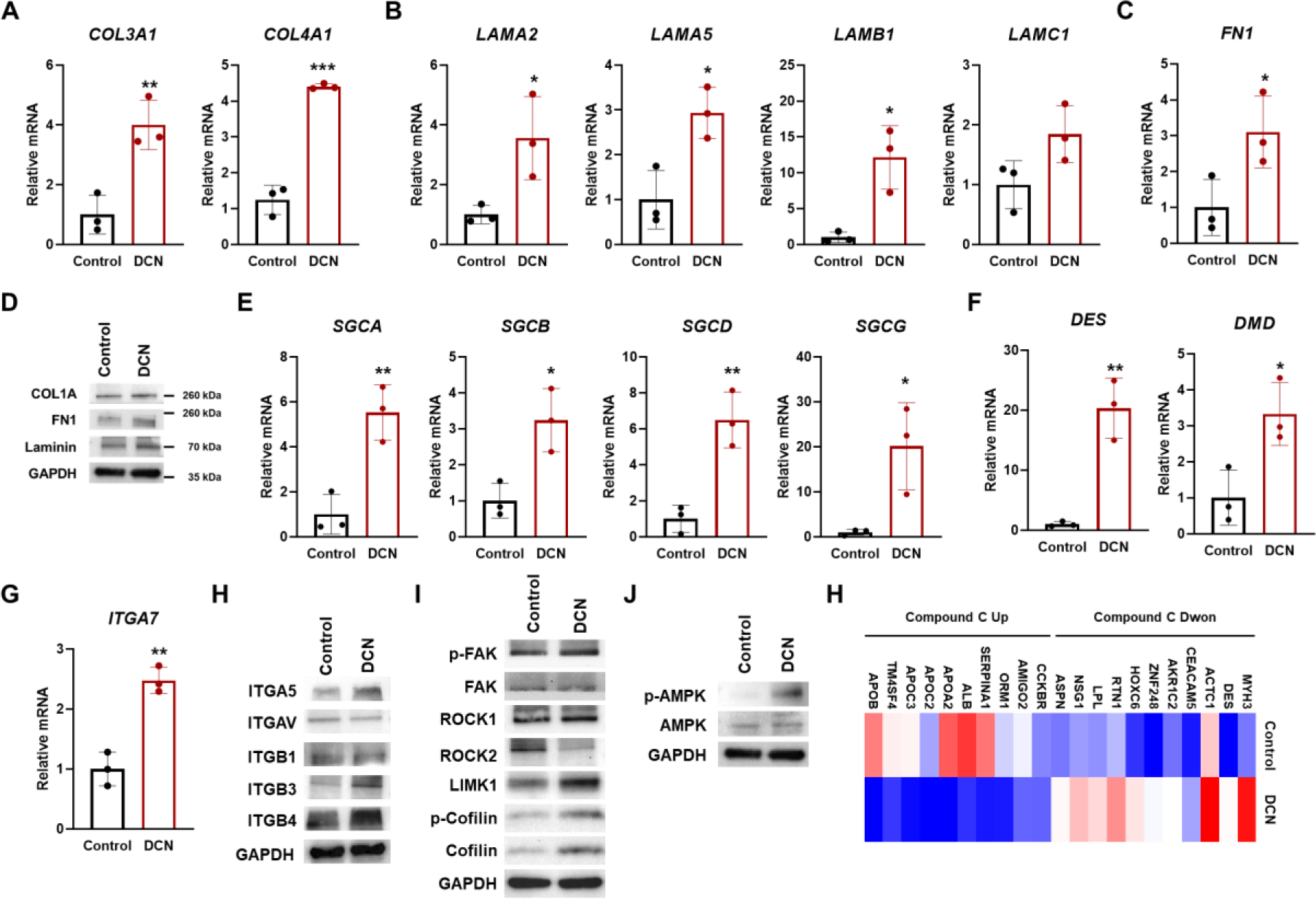
ECM-integrin, calcium signaling, and AMPK signaling pathways are activated by DCN in COs. qRT-PCR of **A,** collagen (*COL3A1* and *COL4A1*), **B,** laminin (*LAMA2*, *LAMA5*, *LAMB1*, and *LAMC1*), and **C,** fibronectin (FN1) markers in control and DCN-treated COs. Values are means ± SDs. n=3 for each group. *p < 0.05, **p < 0.01, ***p < 0.0001. **D,** Western blot analysis of ECM (COL1A1, FN1, and Laminin) markers in control and DCN-treated COs. GAPDH was used as a loading control. qRT-PCR of **E,** sarcoglycan (*SGCA*, *SGCB*, *SGCD*, and *SGCG*), **F,** desmin (*DES*), dystrophin (*DMD*), and **G,** integrin (*ITGA7*) markers in control and DCN-treated COs Values are means ± SDs. n=3 for each group. *p < 0.05, **p < 0.01. Western blot analysis of **H,** Integrin (ITGA5, ITGAV, ITGB1, ITGB3, and ITGB4), **I,** focal adhesion (p-FAK, FAK, ROCK1, ROCK2, LIMK1, p-Cofilin, and Cofilin), and **J,** AMPK signaling (p-AMPK and AMPK) in control and DCN-treated COs. GAPDH was used as a loading control. **K,** LINCS L1000 Chem Pert consensus sigs showing regulated gene expression of up- and down-regulated genes in CC knockdown cells.

MAPK signaling pathway, upregulated by VEGFR2, induces angiogenesis^42^. This study further investigated which aspects of MAPK signaling pathway are primarily responsible for maturation of DCN-treated COs. The protein expression levels of VEGFR2, p-MEK, and p-ERK were higher in DCN-treated COs compared with control (Supplementary Figure 9). However, protein expression levels of p-AKT and p-PI3K were decreased in DCN-treated CO. These results suggest that DCN promotes angiogenesis via upregulating VEGFR2, p-MEK, and p-ERK in COs.

The heart overcomes which is high energy demands predominantly through metabolism of both fatty acid and glucose, which are regulated by AMPK signaling pathway^43^. Protein expression of p-AMPK was significantly higher in DCN-treated COs compared with control (Figure 7J). An increased in mRNA and protein expression of *PGC1A*, *CD36*, and *PPARs*, downstream targets of AMPK, was observed, suggesting that AMPK signaling pathway may be associated with maturation. Moreover, the heatmap showing genes regulated by CC treatment, an AMPK inhibitor, reveals that genes decreased by CC treatment are increased in DCN-treated COs, and genes increased by CC treatment are decreased in DCN-treated COs (Figure 7K). Therefore, these results suggest an association between DCN and AMPK in inducing CO maturation.

### AMPK signaling pathways are activated during maturation of COs

Given the significant activation of AMPK signaling pathway is DCN-treated COs compared with control, we aimed to define roles of AMPK signaling in maturation and angiogenesis of COs. Both control and DCN-treated COs were treated to CC from day 5 to day 30, after which qRT-PCR and western blot analyses were performed. Protein levels of p-AMPK reduced in both CC-treated and DCN and CC-treated COs compared with DCN-treated COs (Figure 8A).

**Figure 8.**
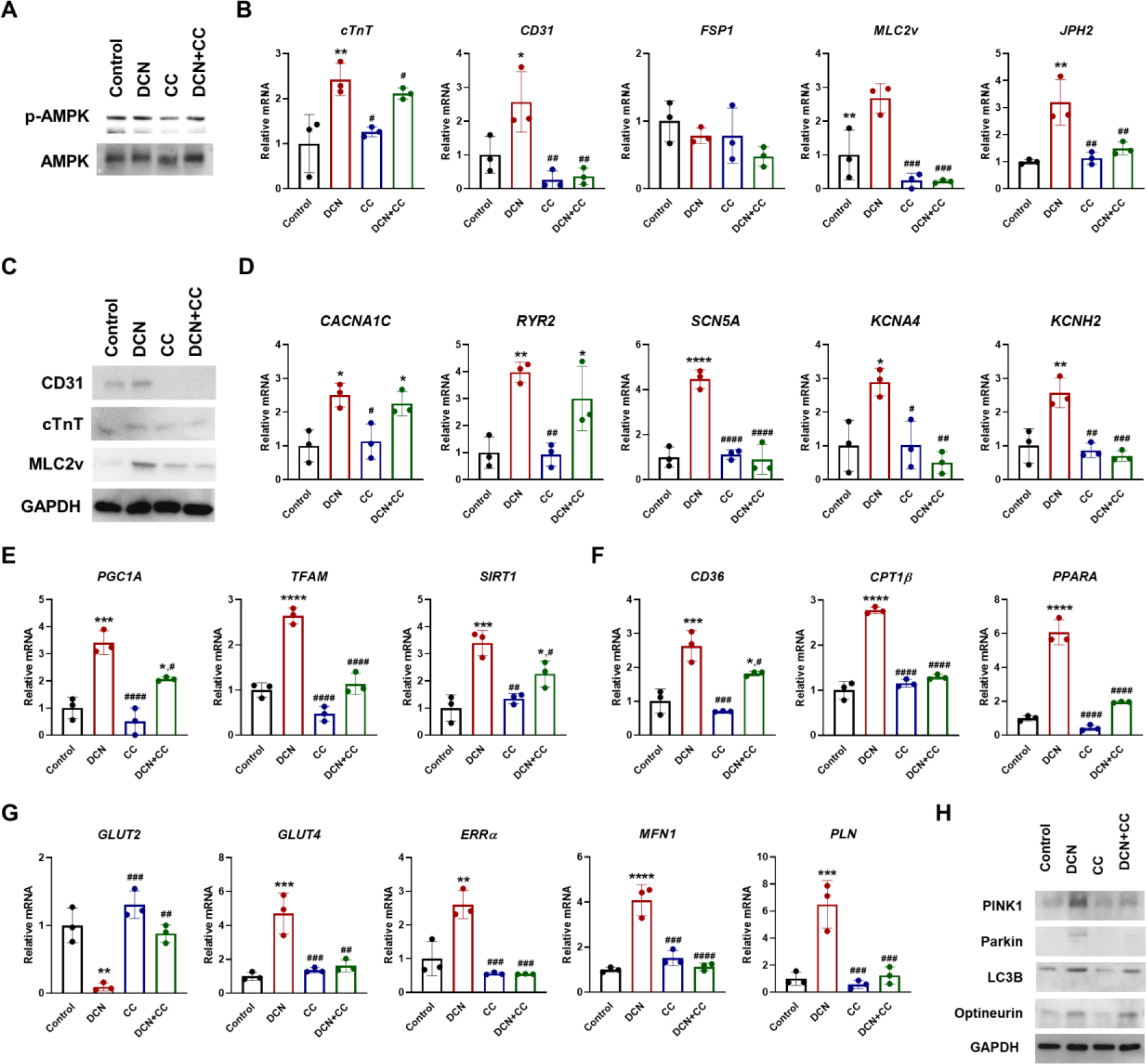
AMPK signaling pathway, activating by DCN, is critical for cardiac maturation and function. **A,** Western blot analysis of AMPK signaling (p-AMPK and AMPK) markers in control, DCN-treated COs, CC-treated COs, and DCN+CC-treated COs. GAPDH was used as a loading control. **B,** qRT-PCR of cardiac component cell (*cTnT*, *CD31*, and *FSP1*) and mature CM (*MLC2v* and *JPH2*) markers in control, DCN-treated COs, CC-treated COs, and DCN+CC-treated COs. Values are means ± SDs. n=3 for each group. *p < 0.05, **p < 0.01 versus control and ##p < 0.01, ###p < 0.001 versus DCN-treated COs. **C,** Western blot analysis of CD31, cTnT, and MLC2v in control, DCN-treated COs, CC-treated COs, and DCN+CC-treated COs. GAPDH was used as a loading control. qRT-PCR of **D,** ion channel (*CACNA1C*, *RYR2*, *SCN5A*, *KCNA4*, and *KCNH2*), **E,** mitochondrial biogenesis (*PGC1A*, *TFAM*, and *SIRT1*), **F,** FAO (*CD36*, *CPT1B*, and *PPARA*), **G,** glycolysis (*GLUT2*, *GLUT4*, *ERRA*, *MFN1*, and *PLN*) markers in control, DCN-treated COs, CC-treated COs, and DCN+CC-treated COs. Values are means ± SDs. n=3 for each group. *p < 0.05, ***p < 0.001, ****p < 0.0001 versus control and #p < 0.05, ##p < 0.01, ####p < 0.0001 versus DCN-treated COs. **H,** Western blot analysis of mitophagy (PINK1, Parkin, LC3B, and Optineurin) markers in control and DCN-treated COs. GAPDH was used as a loading control.

The effects of AMPK knockdown on CM maturation and angiogenesis in DCN-treated COs was examined. mRNA and protein expression levels of *cTnT*, *CD31*, *MLC2v*, and *JPH2* were significantly reduced by AMPK knockdown in both control and DCN-treated COs, as opposed to DCN-treated COs (Figure 8B and 8C). No significant differences in *FSP1* mRNA expression were observed among all groups. These results demonstrated that DCN promotes CM maturation and angiogenesis by modulating AMPK signaling pathway.

The effect of AMPK knockdown on functional maturation in both control and DCN-treated COs was also explored. mRNA expression of sodium channel (*SCN5A*) and potassium channel (*KCNA4* and *KCNH2*) markers was significantly decreased in both CC-treated and DCN and CC-treated COs, compared with DCN-treated COs (Figure 8D). Conversely, calcium channel (*CACNA1C* and *RYR2*) markers showed a decrease in CC-treated COs, with no significant difference observed in DCN and CC-treated COs compared with DCN-treated COs. These results indicate that DCN influences sodium and potassium channel expression through AMPK signaling pathway, whereas calcium channels are not regulated by AMPK signaling pathway.

Moreover, mRNA expression levels of glycolysis (*GLUT4*, *ERRA*, *MFN1*, and *PLN*), FAO (*CD36*, *CPT1B*, and *PPARA*), mitochondrial biogenesis (*PGC1A*, *TFAM*, and *SIRT1*) markers were reduced in both CC-treated and DCN and CC-treated COs compared with DCN-treated COs (Figure 8E-G). Interestingly, protein expression levels of PINK1, Parkin, and LC3B were significantly decreased in CC-treated and DCN and CC-treated COs compared with DCN-treated COs (Figure 8H).

For the first time, we have shown that DCN regulates AMPK signaling pathway and mitophagy, which leads to angiogenesis and functional and metabolic maturation.

## DISCUSSION

This study demonstrated that DCN-treated COs increased and activated mitochondria to produce ATP because COs require many ATPs for continuous contraction–relaxation by spontaneous beating. Therefore, the signaling of mitophagy reportedly increases to reuse aged or damaged mitochondria in cells having increased mitochondrial activity. Mitophagy is an autophagy process that selectively degrades mitochondria that are damaged or stressed by autophagosomes. The mitophagy activity was analyzed using mitochondrial indicator, MitoTracker, and lysosome indicator, LysoTracker, using immunostaining. DCN-treated COs highly expressed mitophagy. These results indicate that DCN-treated COs showed increased mitochondrial number, mitochondrial activity, and mitophagy activity.

AMPK signaling pathway increases mitochondrial biogenesis via activation of PPARs and PGC1A and enhances FAO via CD36 and CPT1B^44^. In DCN-treated COs, upregulation of PPARs and PGC1A led to increased mitochondrial biogenesis, while activation of CD36 and CPT1B enhanced FAO. Therefore, we hypothesized that AMPK plays a crucial role in maturation of COs as a downstream effector of DCN. Interestingly, treatment with CC, an AMPK inhibitor, in DCN-treated COs decreased gene expression of PGC1A, TFAM, and SIRT1, involved in mitochondrial biogenesis. Furthermore, DCN+CC-treated COs reduced expression of CD36, CPT1B, and PPARA, associated with FAO. These results demonstrate that AMPK acts downstream of DCN to regulate mitochondrial content and metabolism.

DCN has been reported to be distributed in mitophagy-associated signaling systems^24,45^. Therefore, mitophagy-associated signaling was analyzed to determine the role of DCN in mitochondrial activity and metabolism in COs. DCN-treated COs highly expressed PINK1 and Parkin. DCN also increased PGC1A expression and induced signaling cascades, including PINK1 and Parkin^46,47^. PINK1 regulates parkin translocation and PINK1-Parkin pathway regulates ubiquitin-dependent mitophagy^47,48^. PINK1-Parkin pathway promotes mitophagy signaling by LC3B and Optineurin^49,50^. These results reveal that DCN regulates mitochondrial activity and mitophagy signaling in COs. Furthermore, treatment with both DCN and CC decreased the expression of PINK1, Parkin, and LC3B, suggesting that AMPK activated by DCN can increase mitophagy activity.

AMPK has been reported to increase glucose uptake via GLUT4 in CMs, which in turn enhances glycolysis^51,52^. Remarkably, expression of GLUT4 was increased in DCN-treated COs but decreased in DCN+CC-treated COs. These results indicate that DCN increases glycolysis in mature CMs via activation of AMPK.

In summary, DCN induced structural, functional, metabolic, and molecular maturation in COs. We demonstrated that AMPK signaling pathway can regulate maturation in DCN-treated COs by controlling mitochondrial biogenesis, FAO, and glycolysis. These mature DCN-treated COs have potential to serve as effective models for drug efficacy testing and disease modeling.

## ACKNOWLEDGMENTS

**Myeong-Hwa Song**: Writing - Original Draft, Methodology, Data curation. **Sungmin Jun**: Investigation. **Seung-Cheol Choi**: Investigation. **Ji Eun Na**: Visualization. **Im Joo Rhyu**: Visualization. **Sun Wook Hwang**: Visualization. **Minji Jeon**: Software. and **Do-Sun Lim**: Supervision

## Nonstandard Abbreviations and Acronyms

CO: Cardiac organoid
CM: Cardiomyocyte
CVD: Cardiovascular disease
CC: Compound C
DCN: Decorin
DEG: differentially expressed gene
EC: Endothelial cell
ECM: Extracellular matrix
FAO: Fatty acid oxidation
FB: Fibroblast
hiPSCs: human induced pluripotent stem cell
hPSC: human pluripotent stem cell
mtDNA: mitochondrial DNA
NGS: normal goat serum
nDNA: nuclear DNA
OCR: Oxygen consumption rate
PFA: paraformaldehyde
PBST: PBS containing 0.1% Tween 20
qRT-PCR: Quantitative reverse transcription polymerase chain reaction
ROS: reactive oxygen species
RT: room temperature
RPMI/B27-Ins: RPMI 1640 medium containing B27 supplement minus insulin
RPMI/B27-VitaA: RPMI 1640 medium with B27 supplement minus vitamin A
t-tubule: Transverse-tubule
TBST: Tris-buffered saline with 0.1% Tween 20

## REFERENCES

1. Zhang D, Pu WT. Exercising engineered heart muscle to maturity. Nature reviews Cardiology. 2018;15:383–384. doi: 10.1038/s41569-018-0032-x

2. Ronaldson-Bouchard K, Ma SP, Yeager K, Chen T, Song L, Sirabella D, Morikawa K, Teles D, Yazawa M, Vunjak-Novakovic G. Advanced maturation of human cardiac tissue grown from pluripotent stem cells. Nature. 2018;556:239–243. doi: 10.1038/s41586-018-0016-3

3. Rossi G, Manfrin A, Lutolf MP. Progress and potential in organoid research. Nature reviews Genetics. 2018;19:671–687. doi: 10.1038/s41576-018-0051-9

4. Matsa E, Burridge PW, Yu KH, Ahrens JH, Termglinchan V, Wu H, Liu C, Shukla P, Sayed N, Churko JM, et al. Transcriptome Profiling of Patient-Specific Human iPSC-Cardiomyocytes Predicts Individual Drug Safety and Efficacy Responses In Vitro. Cell stem cell. 2016;19:311–325. doi: 10.1016/j.stem.2016.07.006

5. Sun C, Kontaridis MI. Physiology of Cardiac Development: From Genetics to Signaling to Therapeutic Strategies. Current opinion in physiology. 2018;1:123–139. doi: 10.1016/j.cophys.2017.09.002

6. Karbassi E, Fenix A, Marchiano S, Muraoka N, Nakamura K, Yang X, Murry CE. Cardiomyocyte maturation: advances in knowledge and implications for regenerative medicine. Nature reviews Cardiology. 2020;17:341–359. doi: 10.1038/s41569-019-0331-x

7. Brade T, Pane LS, Moretti A, Chien KR, Laugwitz KL. Embryonic heart progenitors and cardiogenesis. Cold Spring Harbor perspectives in medicine. 2013;3:a013847. doi: 10.1101/cshperspect.a013847

8. Guo Y, Pu WT. Cardiomyocyte Maturation: New Phase in Development. Circulation research. 2020;126:1086–1106. doi: 10.1161/circresaha.119.315862

9. Leitolis A, Robert AW, Pereira IT, Correa A, Stimamiglio MA. Cardiomyogenesis Modeling Using Pluripotent Stem Cells: The Role of Microenvironmental Signaling. Frontiers in cell and developmental biology. 2019;7:164. doi: 10.3389/fcell.2019.00164

10. Burridge PW, Keller G, Gold JD, Wu JC. Production of de novo cardiomyocytes: human pluripotent stem cell differentiation and direct reprogramming. Cell stem cell. 2012;10:16–28. doi: 10.1016/j.stem.2011.12.013

11. Yang X, Pabon L, Murry CE. Engineering adolescence: maturation of human pluripotent stem cell-derived cardiomyocytes. Circulation research. 2014;114:511–523. doi: 10.1161/circresaha.114.300558

12. Kehat I, Khimovich L, Caspi O, Gepstein A, Shofti R, Arbel G, Huber I, Satin J, Itskovitz-Eldor J, Gepstein L. Electromechanical integration of cardiomyocytes derived from human embryonic stem cells. Nature biotechnology. 2004;22:1282–1289. doi: 10.1038/nbt1014

13. Kojima K, Kaneko T, Yasuda K. Role of the community effect of cardiomyocyte in the entrainment and reestablishment of stable beating rhythms. Biochemical and biophysical research communications. 2006;351:209–215. doi: 10.1016/j.bbrc.2006.10.037

14. Correia C, Koshkin A, Duarte P, Hu D, Teixeira A, Domian I, Serra M, Alves PM. Distinct carbon sources affect structural and functional maturation of cardiomyocytes derived from human pluripotent stem cells. Scientific reports. 2017;7:8590. doi: 10.1038/s41598-017-08713-4

15. Lopaschuk GD, Jaswal JS. Energy metabolic phenotype of the cardiomyocyte during development, differentiation, and postnatal maturation. Journal of cardiovascular pharmacology. 2010;56:130–140. doi: 10.1097/FJC.0b013e3181e74a14

16. Denning C, Borgdorff V, Crutchley J, Firth KS, George V, Kalra S, Kondrashov A, Hoang MD, Mosqueira D, Patel A, et al. Cardiomyocytes from human pluripotent stem cells: From laboratory curiosity to industrial biomedical platform. Biochimica et biophysica acta. 2016;1863:1728–1748. doi: 10.1016/j.bbamcr.2015.10.014

17. Richards DJ, Coyle RC, Tan Y, Jia J, Wong K, Toomer K, Menick DR, Mei Y. Inspiration from heart development: Biomimetic development of functional human cardiac organoids. Biomaterials. 2017;142:112–123. doi: 10.1016/j.biomaterials.2017.07.021

18. Günthel M, Barnett P, Christoffels VM. Development, Proliferation, and Growth of the Mammalian Heart. Molecular therapy: the journal of the American Society of Gene Therapy. 2018;26:1599–1609. doi: 10.1016/j.ymthe.2018.05.022

19. Nashimoto Y, Hayashi T, Kunita I, Nakamasu A, Torisawa YS, Nakayama M, Takigawa-Imamura H, Kotera H, Nishiyama K, Miura T, et al. Integrating perfusable vascular networks with a three-dimensional tissue in a microfluidic device. Integrative biology: quantitative biosciences from nano to macro. 2017;9:506–518. doi: 10.1039/c7ib00024c

20. Zhao X, Xu Z, Xiao L, Shi T, Xiao H, Wang Y, Li Y, Xue F, Zeng W. Review on the Vascularization of Organoids and Organoids-on-a-Chip. Frontiers in bioengineering and biotechnology. 2021;9:637048. doi: 10.3389/fbioe.2021.637048

21. Järvinen TA, Prince S. Decorin: A Growth Factor Antagonist for Tumor Growth Inhibition. BioMed research international. 2015;2015:654765. doi: 10.1155/2015/654765

22. Gubbiotti MA, Buraschi S, Kapoor A, Iozzo RV. Proteoglycan signaling in tumor angiogenesis and endothelial cell autophagy. Seminars in cancer biology. 2020;62:1–8. doi: 10.1016/j.semcancer.2019.05.003

23. Vu TT, Marquez J, Le LT, Nguyen ATT, Kim HK, Han J. The role of decorin in cardiovascular diseases: more than just a decoration. Free radical research. 2018;52:1210–1219. doi: 10.1080/10715762.2018.1516285

24. Neill T, Torres A, Buraschi S, Owens RT, Hoek JB, Baffa R, Iozzo RV. Decorin induces mitophagy in breast carcinoma cells via peroxisome proliferator-activated receptor γ coactivator-1α (PGC-1α) and mitostatin. The Journal of biological chemistry. 2014;289:4952–4968. doi: 10.1074/jbc.M113.512566

25. Iozzo RV, Buraschi S, Genua M, Xu SQ, Solomides CC, Peiper SC, Gomella LG, Owens RC, Morrione A. Decorin antagonizes IGF receptor I (IGF-IR) function by interfering with IGF-IR activity and attenuating downstream signaling. The Journal of biological chemistry. 2011;286:34712–34721. doi: 10.1074/jbc.M111.262766

26. Sorensen DW, van Berlo JH. The Role of TGF-β Signaling in Cardiomyocyte Proliferation. Current heart failure reports. 2020;17:225–233. doi: 10.1007/s11897-020-00470-2

27. Chisalita SI, Johansson GS, Liefvendahl E, Bäck K, Arnqvist HJ. Human aortic smooth muscle cells are insulin resistant at the receptor level but sensitive to IGF1 and IGF2. Journal of molecular endocrinology. 2009;43:231–239. doi: 10.1677/jme-09-0021

28. Vučković S, Dinani R, Nollet EE, Kuster DWD, Buikema JW, Houtkooper RH, Nabben M, van der Velden J, Goversen B. Characterization of cardiac metabolism in iPSC-derived cardiomyocytes: lessons from maturation and disease modeling. Stem cell research & therapy. 2022;13:332. doi: 10.1186/s13287-022-03021-9

29. Sun X, Nunes SS. Bioengineering Approaches to Mature Human Pluripotent Stem Cell-Derived Cardiomyocytes. Frontiers in cell and developmental biology. 2017;5:19. doi: 10.3389/fcell.2017.00019

30. Hong T, Shaw RM. Cardiac T-Tubule Microanatomy and Function. Physiological reviews. 2017;97:227-252. doi: 10.1152/physrev.00037.2015

31. Wu P, Deng G, Sai X, Guo H, Huang H, Zhu P. Maturation strategies and limitations of induced pluripotent stem cell-derived cardiomyocytes. Bioscience reports. 2021;41. doi: 10.1042/bsr20200833

32. Lascano EC, Negroni JA. Gap junctions in preconditioning against arrhythmias. Cardiovascular research. 2007;74:341–342. doi: 10.1016/j.cardiores.2007.04.003

33. Bers DM. Cardiac excitation-contraction coupling. Nature. 2002;415:198-205. doi: 10.1038/415198a

34. Keung W, Boheler KR, Li RA. Developmental cues for the maturation of metabolic, electrophysiological and calcium handling properties of human pluripotent stem cell-derived cardiomyocytes. Stem cell research & therapy. 2014;5:17. doi: 10.1186/scrt406

35. Jiang Y, Park P, Hong SM, Ban K. Maturation of Cardiomyocytes Derived from Human Pluripotent Stem Cells: Current Strategies and Limitations. Molecules and cells. 2018;41:613–621. doi: 10.14348/molcells.2018.0143

36. Mongirdienė A, Skrodenis L, Varoneckaitė L, Mierkytė G, Gerulis J. Reactive Oxygen Species Induced Pathways in Heart Failure Pathogenesis and Potential Therapeutic Strategies. Biomedicines. 2022;10. doi: 10.3390/biomedicines10030602

37. Momtahan N, Crosby CO, Zoldan J. The Role of Reactive Oxygen Species in In Vitro Cardiac Maturation. Trends in molecular medicine. 2019;25:482–493. doi: 10.1016/j.molmed.2019.04.005

38. Grynberg A, Demaison L. Fatty acid oxidation in the heart. Journal of cardiovascular pharmacology. 1996;28 Suppl 1:S11–17. doi: 10.1097/00005344-199600003-00003

39. Garbern JC, Lee RT. Mitochondria and metabolic transitions in cardiomyocytes: lessons from development for stem cell-derived cardiomyocytes. Stem cell research & therapy. 2021;12:177. doi: 10.1186/s13287-021-02252-6

40. Villarejo-Zori B, Jiménez-Loygorri JI, Boya P. HIF1α or mitophagy: which drives cardiomyocyte differentiation? Cell stress. 2020;4:95–98. doi: 10.15698/cst2020.05.219

41. Dorn GW, 2nd. Parkin-dependent mitophagy in the heart. Journal of molecular and cellular cardiology. 2016;95:42–49. doi: 10.1016/j.yjmcc.2015.11.023

42. Beamer B, Hettrich C, Lane J. Vascular endothelial growth factor: an essential component of angiogenesis and fracture healing. HSS journal: the musculoskeletal journal of Hospital for Special Surgery. 2010;6:85–94. doi: 10.1007/s11420-009-9129-4

43. Dyck JR, Lopaschuk GD. AMPK alterations in cardiac physiology and pathology: enemy or ally? The Journal of physiology. 2006;574:95–112. doi: 10.1113/jphysiol.2006.109389

44. Sarikhani M, Garbern JC, Ma S, Sereda R, Conde J, Krähenbühl G, Escalante GO, Ahmed A, Buenrostro JD, Lee RT. Sustained Activation of AMPK Enhances Differentiation of Human iPSC-Derived Cardiomyocytes via Sirtuin Activation. Stem cell reports. 2020;15:498–514. doi: 10.1016/j.stemcr.2020.06.012

45. Chen G, Kroemer G, Kepp O. Mitophagy: An Emerging Role in Aging and Age-Associated Diseases. Frontiers in cell and developmental biology. 2020;8:200. doi: 10.3389/fcell.2020.00200

46. Peng K, Xiao J, Yang L, Ye F, Cao J, Sai Y. Mutual Antagonism of PINK1/Parkin and PGC-1α Contributes to Maintenance of Mitochondrial Homeostasis in Rotenone-Induced Neurotoxicity. Neurotoxicity research. 2019;35:331–343. doi: 10.1007/s12640-018-9957-4

47. Quinn PMJ, Moreira PI, Ambrósio AF, Alves CH. PINK1/PARKIN signalling in neurodegeneration and neuroinflammation. Acta neuropathologica communications. 2020;8:189. doi: 10.1186/s40478-020-01062-w

48. Ge P, Dawson VL, Dawson TM. PINK1 and Parkin mitochondrial quality control: a source of regional vulnerability in Parkinson’s disease. Molecular neurodegeneration. 2020;15:20. doi: 10.1186/s13024-020-00367-7

49. Vincow ES, Merrihew G, Thomas RE, Shulman NJ, Beyer RP, MacCoss MJ, Pallanck LJ. The PINK1-Parkin pathway promotes both mitophagy and selective respiratory chain turnover in vivo. Proceedings of the National Academy of Sciences of the United States of America. 2013;110:6400–6405. doi: 10.1073/pnas.1221132110

50. Lin Q, Li S, Jiang N, Shao X, Zhang M, Jin H, Zhang Z, Shen J, Zhou Y, Zhou W, et al. PINK1-parkin pathway of mitophagy protects against contrast-induced acute kidney injury via decreasing mitochondrial ROS and NLRP3 inflammasome activation. Redox biology. 2019;26:101254. doi: 10.1016/j.redox.2019.101254

51. Russell RR, 3rd, Li J, Coven DL, Pypaert M, Zechner C, Palmeri M, Giordano FJ, Mu J, Birnbaum MJ, Young LH. AMP-activated protein kinase mediates ischemic glucose uptake and prevents postischemic cardiac dysfunction, apoptosis, and injury. The Journal of clinical investigation. 2004;114:495–503. doi: 10.1172/jci19297

52. Ji L, Zhang X, Liu W, Huang Q, Yang W, Fu F, Ma H, Su H, Wang H, Wang J, et al. AMPK-regulated and Akt-dependent enhancement of glucose uptake is essential in ischemic preconditioning-alleviated reperfusion injury. PloS one. 2013;8:e69910. doi: 10.1371/journal.pone.0069910

